# Hepatic Mitochondrial Remodeling is Mechanistically Linked to Insulin Resistance in Nonalcoholic Fatty Liver Disease

**DOI:** 10.1101/2020.01.02.892992

**Authors:** Chris E. Shannon, Mukundan Ragavan, Juan Pablo Palavicini, Marcel Fourcaudot, Terry Bakewell, Eunsook S. Jin, Muniswamy Madesh, Craig R. Malloy, Xianlin Han, Matthew E. Merritt, Luke Norton

## Abstract

Insulin resistance and altered hepatic mitochondrial function are central features of type 2 diabetes (T2D) and non-alcoholic fatty liver disease (NAFLD), but the etiological role of these processes in disease progression remains unclear. We investigated the molecular links between insulin resistance, mitochondrial remodeling, and hepatic lipid accumulation in a rodent model of T2D / NAFLD. Livers from obese, insulin resistant mice displayed augmented mitochondrial content and increased TCA cycle and pyruvate dehydrogenase (PDH) activities. Insulin sensitization with pioglitazone mitigated pyruvate-driven TCA cycle activity and PDH activation via both covalent (PDK4 and PDP2) and allosteric (intracellular pyruvate availability) mechanisms. Interestingly, improvements in insulin sensitivity and mitochondrial function were entirely dissociated from changes in hepatic triglycerides, diacylglycerides or fatty acids. Instead, we show that the mitochondrial phospholipid cardiolipin undergoes pathological remodeling in livers from obese mice and that this is reversed by insulin sensitization. Our findings identify targetable mitochondrial features of T2D and NAFLD and highlight the benefit of insulin sensitization in managing the clinical burden of obesity-associated disease.

## INTRODUCTION

Type 2 diabetes mellitus (T2D) increases the risk of developing non-alcoholic fatty liver disease (NAFLD), and NAFLD patients with T2D are more likely to progress towards the more severe maladies of non-alcoholic steatohepatitis (NASH) and hepatocellular carcinoma (1, 2). Although the precise events linking T2D and NAFLD remain unclear, their shared association with obesity and insulin resistance suggests a common underlying pathology characterized by ectopic lipid accumulation and elevated rates of hepatic glucose production (3-5).

Studies in humans have identified alterations in hepatic mitochondrial metabolism as a central feature of T2D and NAFLD (5, 6), including the remodeling of mitochondrial lipids (7), increases in mitochondrial mass and respiratory capacity (8), altered acetyl-CoA metabolism (9) and excessive tricarboxylic acid (TCA) cycle turnover (5). By driving sustained elevations in oxidative stress, these adaptations are believed to contribute to the inflammatory, fibrotic milieu associated with disease progression (8) and warrant the development of therapies specifically targeting hepatic mitochondrial metabolism.

Pioglitazone is an insulin-sensitizing thiazolidinedione prescribed for the treatment of T2D (10) and has shown considerable efficacy in improving NAFLD and NASH outcomes (11, 12). Although typically ascribed to adipose tissue PPAR-γ-mediated reductions in systemic lipid concentrations and ectopic lipid accumulation (13-15), the disease-modifying effects of pioglitazone may also involve direct effects on liver metabolism. For example, pioglitazone acutely suppresses glucose production in cultured hepatocytes (16, 17) and perfused livers (18) and was recently found to decrease liver TCA cycle flux in a mouse model of NASH (19). The molecular mechanisms responsible for modulation of hepatic mitochondrial metabolism by pioglitazone are uncertain but data from our lab (16) and others (20) indicate that they may involve the inhibition of mitochondrial pyruvate fluxes.

Here, we investigated hepatic mitochondrial adaptation in a rodent model of liver fat accumulation and insulin resistance, and under conditions where insulin sensitivity was improved with pioglitazone, using a combination of ^2^H/^13^C NMR metabolic flux analyses, high-resolution shotgun lipidomics, and molecular approaches.

### RESULTS

### Hepatic mitochondrial content parallels liver insulin resistance in obese mice

To investigate the relationship between insulin resistance and hepatic mitochondrial adaptation, we first compared protein markers of mitochondrial content from the livers of 16-week old wild-type control mice (WT) with those from aged-matched obese (ob/ob) mice (OB-CON) and obese mice fed a diet supplemented with the insulin sensitizer pioglitazone (300 mg/kg diet) for four weeks (OB-PIO). Cytochrome C oxidase IV, a subunit from complex IV of the respiratory transport chain, was increased in OB-CON mice, whereas the highly abundant outer mitochondrial membrane protein VDAC1 was unchanged (Figure 1A and B). We next determined citrate synthase activity in liver homogenates as a more specific measure of hepatic mitochondrial content.

**Figure 1:**
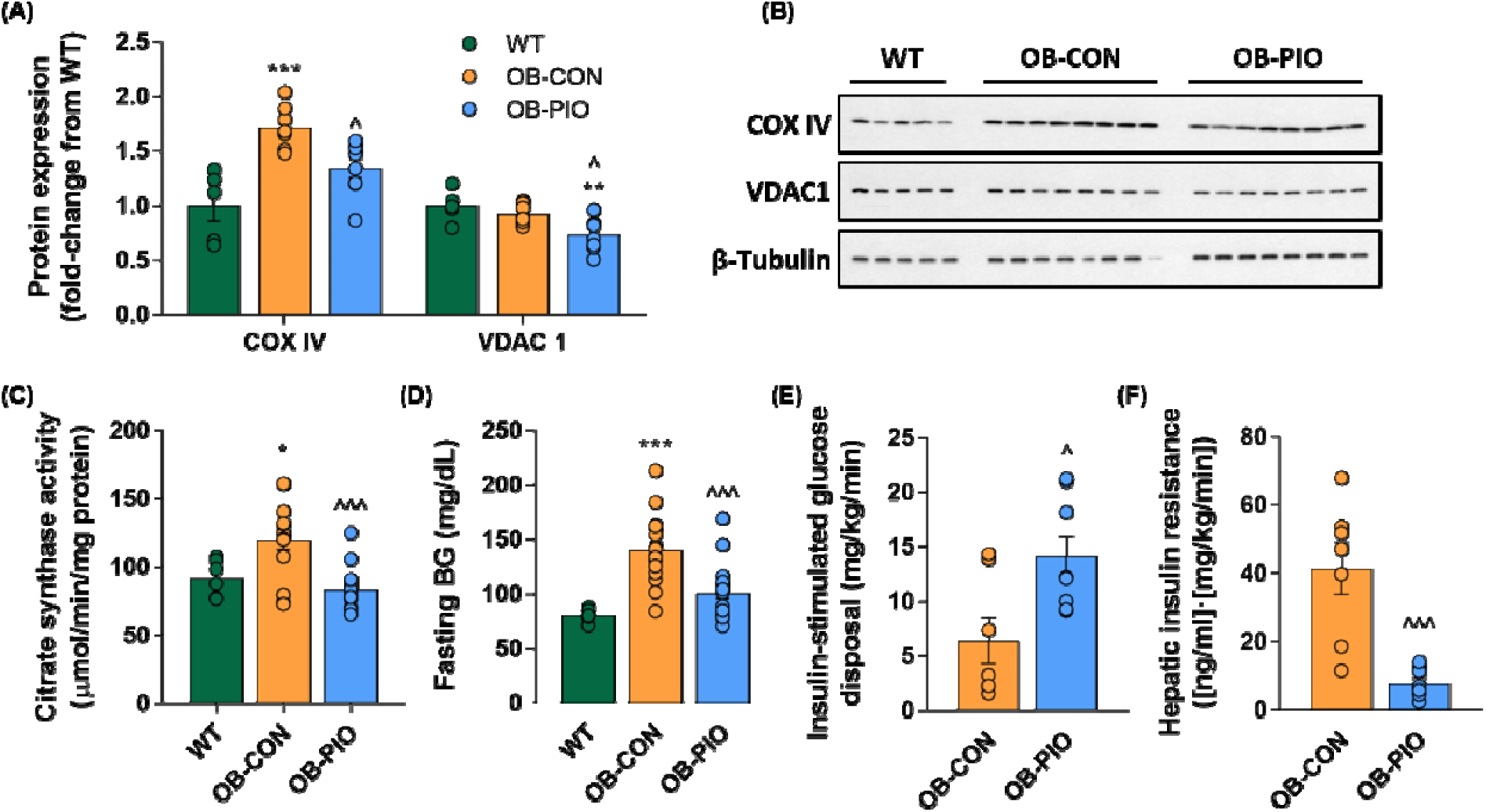
Hepatic mitochondrial content parallels liver insulin resistance in obese mice: (A) Liver protein expression (relative to wild-type mice) and (B) representative western blots of mitochondrial markers cytochrome C oxidase IV and VDAC1, normalized to β–Tubulin control, in livers from wild-type, untreated obese mice (OB-CON) and obese mice treated with pioglitazone (∼20 mg/kg/day) for four weeks (OB-PIO). (C) Hepatic citrate synthase activity measured spectrophotometrically and (D) fasting blood glucose in WT, OB-CON and OB-PIO mice. (E) Whole-body insulin sensitivity assessed during euglycemic-hyperinsulinemic clamp and (F) hepatic insulin resistance index (fasting plasma insulin multiplied by basal endogenous glucose production) in OB-CON and OB-PIO mice. Data are mean ± SE for at least five mice per group, as indicated on individual figure panels. *P<0.05, **P<0.01, ***P<0.001 vs WT; ^P<0.05, ^^^P<0.001 vs OB-CON.

Citrate synthase activity was 30% greater in OB-CON compared to WT mice (Figure 1C). These changes are in accordance with the excessive oxidative liver metabolism and mitochondrial content reported in other models of insulin resistance and NAFLD (9, 21). Remarkably, insulin sensitization with pioglitazone reduced all markers of mitochondrial content measured (Figures 1A-C). This normalization of mitochondrial content was paralleled by improvements in glucose homeostasis (Figure 1D and Supplemental Figure 1), as well as in whole-body (Figure 1E) and liver-specific (Figure 1F) insulin sensitivity. Consistent with human studies (10, 22), this insulin sensitization occurred despite significantly greater weight gain and fat mass in pioglitazone-treated mice (Supplemental Table 1).

### Insulin sensitization suppresses hepatic mitochondrial pyruvate metabolism

We next explored whether alterations in mitochondrial pyruvate metabolism were involved in the restoration of glucose homeostasis by pioglitazone. Overnight fasted OB-CON and OB-PIO mice were administered intraperitoneally with a bolus injection of [2,3-^13^C]pyruvate for the assessment of mitochondrial pyruvate flux into glucose (labelling scheme depicted in Figure 2A). NMR analysis of the resultant ^13^C isotopomers in plasma glucose revealed production of [1,2-^13^C] and [5,6-^13^C]glucose in fasted OB-CON and OB-PIO mice (Supplemental Figure 2A), reflecting direct pyruvate-driven gluconeogenesis (23). However, we also observed enrichment of the C-3 and C-4 glucose carbons, demonstrating extensive TCA cycle turnover, in OB-CON but not OB-PIO mice (Figure 2B). Matching simulated spectra from tcaSIM (24) supported a model in which TCA cycle flux was slowed by pioglitazone treatment (Figure 2C).

**Figure 2:**
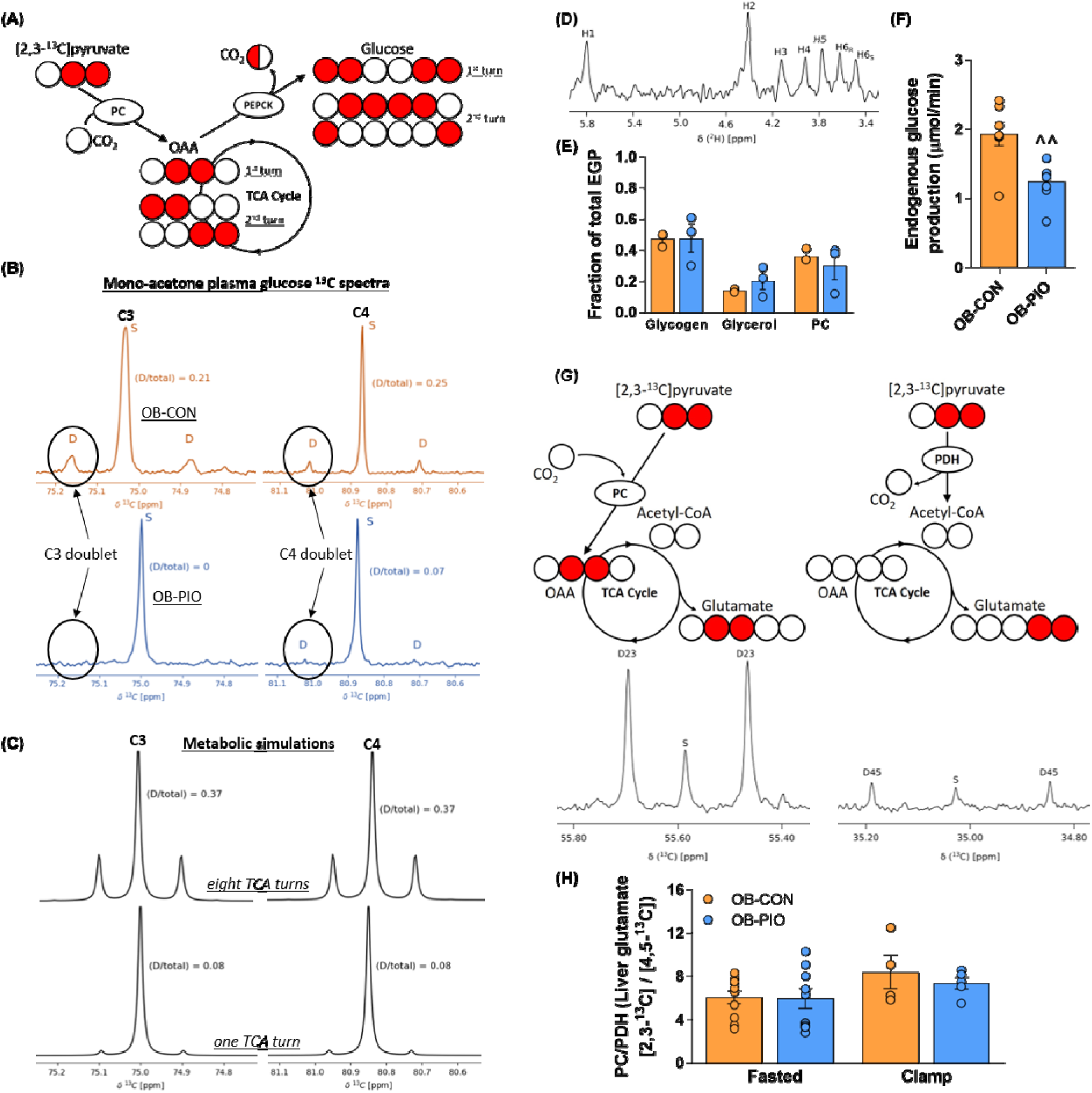
Insulin sensitization suppresses mitochondrial pyruvate metabolism in liver: (A) Schematic illustrating the labelling pattern of plasma glucose following administration of [2,3-^13^C] pyruvate. PC-dependent gluconeogenesis generates predominantly [1,2-^13^C] and [5,6-^13^C] glucose isotopomers, but carbon scrambling through excessive TCA cycle turnover can yield enrichment at interior glucose carbons (i.e. [2,3-^13^C] and [4,5-^13^C] isotopomers). PC pyruvate carboxylase, PEPCK pyruvate carboxykinase. (B) ^13^C spectra showing increased C-3 (left) and C-4 (right) resonance of mono-acetone glucose synthesized from plasma in OB-CON (top) compared to OB-PIO (bottom) mice. (C) Output of tcaSIM metabolic simulations for C3 (left) and C4 (right) run for eight turns (top) and one turn (bottom) of the TCA cycle using lactate enrichments obtained experimentally. (D) Representative mono-acetone glucose ^2^H NMR spectra derived from plasma glucose of fasted OB-CON and OB-PIO mice. (E) Fractional contribution from glycogenolysis (1- [H5/H2]), glycerol-driven gluconeogenesis ([H5-H6S]/H2) and PC-driven gluconeogenesis (H6s/H2) calculated from mono-acetone glucose 2H NMR spectra (n=3). (F) Endogenous glucose production in fasted OB-CON (n=7) and OB-PIO (n=9) mice measured using 3-3H glucose. (G) Schematic (top) showing determination of the ratio between pyruvate carboxylase (PC) and pyruvate dehydrogenase (PDH) flux from the labelling pattern of liver glutamate following [2,3-13C] pyruvate metabolism through either PC (left panel, yielding [2,3-13C] glutamate) or PDH (right panel, yielding [4,5-13C] glutamate) and representative glutamate 13C NMR spectra (bottom). (H) Quantified ratio of liver PC/PDH, reflecting relative mitochondrial anaplerosis / oxidative fluxes determined in OB-CON and OB-PIO mice in basal state and following

The coupling between TCA cycle turnover and glucose production is mediated by pyruvate carboxylase (PC) flux (9) and, as such, we determined the contribution of PC flux to endogenous glucose production (EGP) in parallel experiments using ^2^H_2_O (Figure 2D). Gluconeogenesis (GNG) from PC-flux accounted for a similar proportion of total EGP in overnight fasted OB-CON and OB-PIO mice (Figure 2E). However total EGP, assessed by 3-^3^Hglucose, was 35% lower in OB-PIO mice compared to OB-CON (Figure 2F). Scaling to absolute EGP suggested that PC-driven GNG was double in OB-CON compared to OB-PIO mice (∼0.70 vs 0.37 µmol/min) and demonstrate that hepatic insulin sensitization suppresses excessive mitochondrial metabolism and restricts GNG from the level of the TCA cycle.

As PC-driven glucose production was reduced by pioglitazone, we postulated that alternate pathways of mitochondrial pyruvate disposal might also be affected, such as pyruvate dehydrogenase (PDH). Following [2,3-^13^C]pyruvate administration in fasted OB-CON and OB-PIO mice, we measured the ratio of [2,3-^13^C] / [4,5-^13^C]glutamate in *in situ* freeze-clamped livers, reflecting the entry of pyruvate into the TCA cycle via PC versus PDH (Figure 2G). Unexpectedly, the PC/PDH ratio was unchanged by pioglitazone treatment (Figure 2H). As the hepatic PC/PDH ratio may be sensitive to the fasting-fed transition (23), we repeated these experiments in OB-CON and OB-PIO mice following euglycemic-hyperinsulinemic clamps. Despite a 63% reduction in EGP in OB-PIO compared to OB-CON under insulin-stimulated conditions (Supplemental Figure 2B), the PC/PDH ratio remained unchanged (Figure 2H).

### Pyruvate dehydrogenase activity is increased in obese livers and reduced by insulin sensitization

The constancy of hepatic PC/PDH despite reduced PC flux implied that PDH flux was also lower in OB-PIO mice, which prompted us to investigate the role of PDH in hepatic insulin resistance. Interestingly, we found that protein levels of the PDH catalytic subunit E1α were increased by 70% in OB-CON mice compared to WT and remained elevated in OB-PIO mice (Figure 3A). However, the enzymatic activity of PDH is primarily controlled by its catalytic activation status, which we subsequently measured in *in situ* freeze-clamped livers. Strikingly, and consistent with the increased E1α protein levels, basal PDH activation status was 6-fold higher in OB-CON mice compared to WT but was partially normalized in OB-PIO mice (Figure 3B). This blunting of hepatic PDH activity by pioglitazone was corroborated in mice undergoing insulin clamp studies (Figure 3C). Thus, increased PDH activity may contribute to the mitochondrial dysfunction present in livers from obese mice and is abated by insulin sensitization.

**Figure 3:**
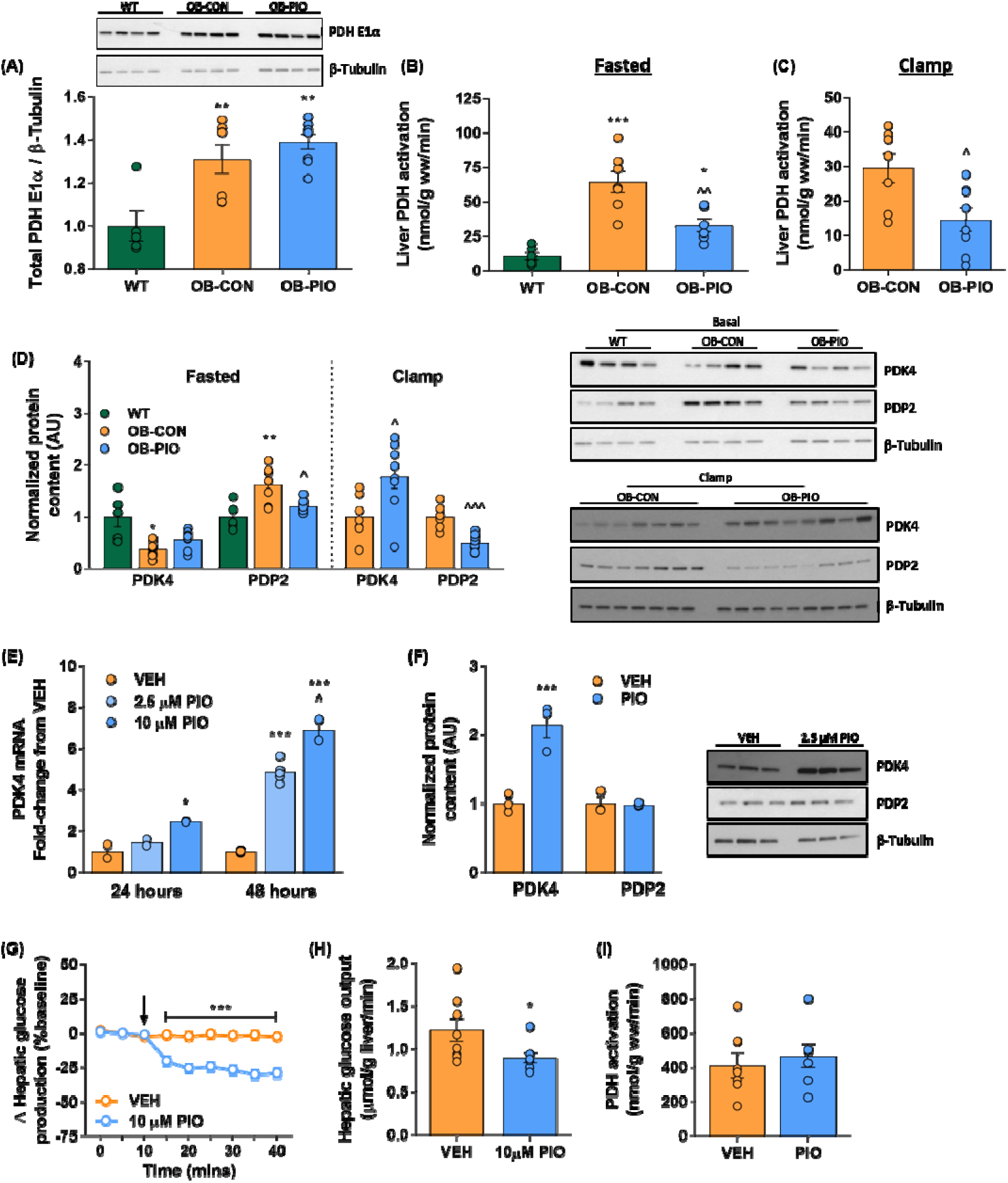
Pyruvate dehydrogenase activity is increased in obese livers and reduced by an insulin sensitizer: (A) Protein expression of the E1α catalytic subunit of the pyruvate dehydrogenase complex (normalized to β-tubulin control), normalized to wild-type levels, in fasted WT, OB-CON and OB-PIO mice, with representative blots. (B) Liver PDH activation measured in *in situ* freeze-clamped livers from overnight fasted OB-CON and OB-PIO mice. (C) Liver PDH activation in OB-CON and OB-PIO mice following insulin clamp. (D) Liver pyruvate dehydrogenase kinase 4 (PDK4) and pyruvate dehydrogenase phosphatase 2 (PDP2) protein expression in fasted WT, OB-CON and OB-PIO and in OB-CON and OB-PIO mice following insulin clamp studies. (E) PDK4 mRNA in primary hepatocytes from treated with DMSO or 2.5 – 10 µM pioglitazone for 24-48 hours. (F) PDK4 and PDP2 protein expression in hepatocytes isolated from wild-type mice treated with DMSO or 2.5 µM pioglitazone for 48 hours, with representative western blot images. (G) Hepatic glucose output expressed as % change from baseline from wild-type livers perfused with buffer containing pyruvate / lactate (1 / 10 mM). Pioglitazone (final concentration 10 µM) or vehicle (DMSO) were added to the perfusion after 10 minutes. (H) Steady-state hepatic glucose output over the final 10 minutes of liver perfusion with vehicle or 10 µM pioglitazone. (I) Hepatic PDH activation determined in in situ freeze-clamped isolated livers following 40 minutes of perfusion under conditions described above. *P<0.05, **P<0.01, P<0.001 vs WT (or vs VEH in panels G-H); ^P<0.05, ^^^P<0.001 vs OB-CON (or vs 2.5µM PIO in panel E).

The activity of PDH is under covalent regulation by the competing actions of the PDH inhibitory kinases (PDKs) and activating phosphatases (PDPs), with PDK4 and PDP2 being the isoforms of primary importance in murine liver (25, 26). Under fasted conditions, hepatic PDK4 protein expression was reduced in OB-CON, but not OB-PIO, compared to WT mice (Figure 3D). Conversely, PDP2 protein levels were increased by 60% in OB-CON and completely normalized in OB-PIO (Figure 3D). Moreover, following insulin clamp studies, PDK4 was 78% greater, whereas PDP2 expression was 50% lower, in OB-PIO compared to OB-CON mice (Figure 3D). Thus, reduced PDK4 and increased PDP2 expression *in vivo* mechanistically contributes to the excessive PDH activation and mitochondrial dysfunction in livers from obese mice, whilst insulin sensitization reverses these changes to suppress PDH activity. Hepatic PDK4 and PDP2 may be responsive to circulating concentrations of free fatty acids and insulin (27).

Since plasma insulin concentrations were lower in OB-PIO mice (Table 1), we next investigated if pioglitazone influenced PDK4 and PDP2 expression independent of insulin, in isolated hepatocytes from wild-type mice. Treatment of primary hepatocytes with pioglitazone for 24-48 hours resulted in a dose- and time-dependent increase in PDK4 mRNA (Figure 3E) which translated into a 2-fold increase in PDK4 protein after 48 hours (Figure 3F). However, no effects of pioglitazone treatment on PDP2 protein expression were observed (Figure 3F), suggesting that pioglitazone-mediated alterations in PDK4 and PDP2 may occur through divergent mechanisms. These experiments in isolated hepatocytes also suggested that pioglitazone may modulate hepatic PDH activation directly, in a liver-autonomous manner. To explore this possibility, we conducted experiments in perfused livers from untreated wild-type mice. Hepatic glucose output was stable over the 40-minute measurement period under control conditions, whereas infusion of 10 µM pioglitazone caused a ∼30% reduction in glucose production rates (Figure 3G-H). However, PDH activation was no different between control and pioglitazone-perfused livers (Figure 3I). Thus, whereas the inhibition of hepatic glucose output by pioglitazone occurs rapidly (within minutes) in isolated perfused livers, the suppression of hepatic PDH activation may require more prolonged (i.e. 24 hours) pioglitazone exposure.

### Hepatic acetyl-CoA concentrations are diminished in obese mice

In addition to post-translational control by PDK4 and PDP2, PDH is allosterically activated by a reduction in the ratio of acetyl-CoA to free coenzyme A (Acetyl-CoA / CoASH) (28). Since hepatic acetyl-CoA metabolism may be perturbed under insulin-resistant conditions (9), we measured acetyl-CoA and CoASH concentrations in *in situ* freeze-clamped livers using a sensitive radioenzymatic assay (29). Acetyl-CoA (Figure 4A) and CoASH (Figure 4B) were decreased by 9-fold and 2-fold, respectively, in OB-CON compared to WT mice, resulting in a 3-fold reduction in acetyl-CoA / CoASH (Figure 4C). We also determined acetylcarnitine concentrations, which parallel changes in the mitochondrial acetyl-CoA pool via the carnitine acetyltransferase reaction (30). Acetylcarnitine was 5-fold lower in livers from OB-CON versus WT mice (Figure 4D), providing independent validation of our measurement of acetyl-CoA. This depletion of hepatic acetyl-CoA may thus contribute to the excessive PDH activity in livers from obese mice and is consistent with increased oxidative flux through the TCA cycle as observed in humans with NAFLD (9). We also asked whether the restoration of hepatic acetyl-CoA levels could explain the lower PDH activity in pioglitazone-treated mice. However, insulin sensitization with pioglitazone had no impact upon hepatic acetyl-CoA concentrations, acetyl-CoA / CoASH or acetylcarnitine concentrations (Figure 4A-D).

**Figure 4:**
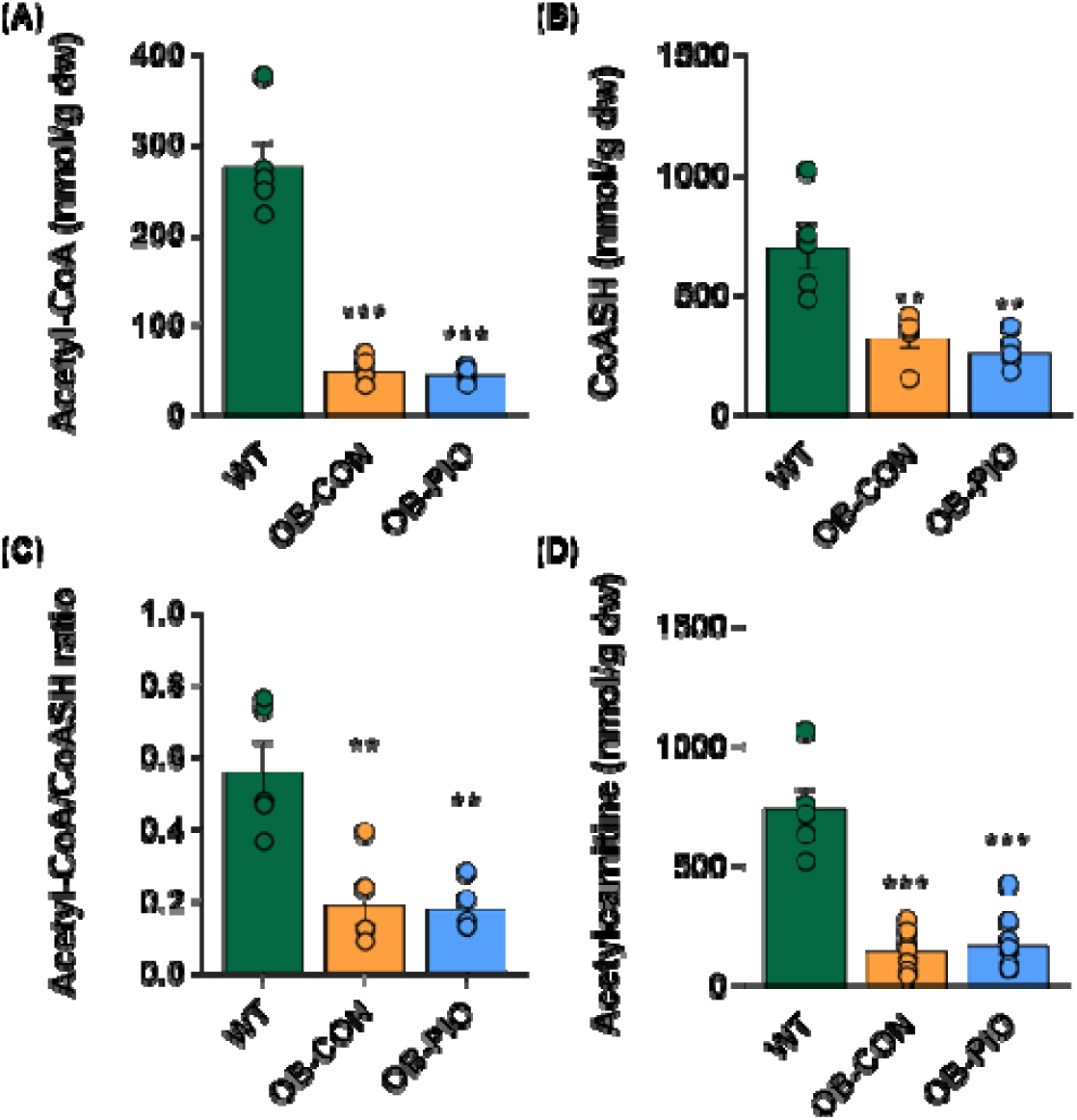
Acetyl-CoA is depleted in obese livers: (A) Acetyl-CoA, (B) free coenzyme A, (C) acetyl-CoA / CoASH ratio and (D) acetylcarnitine concentrations in livers from overnight-fasted wild-type (WT), untreated obese (OB-CON) and pioglitazone-treated obese (OB-PIO) mice. **P<0.01, ***P<0.001 vs WT.

### Insulin sensitization is associated with a reduction in glycogenolysis and hepatic pyruvate availability

Alterations in hepatic acetyl-CoA disposal have previously been attributed to increased pyruvate availability in obese individuals (9). Consistent with this, hepatic pyruvate concentrations were increased by 2.5-fold in OB-CON compared to WT mice and were completely normalized by pioglitazone treatment (Figure 5A). Thus, changes in intracellular pyruvate availability, an allosteric activator of PDH (31), may represent an important mechanism linking insulin resistance with hepatic mitochondrial function. Hepatic pyruvate availability is determined by the balance between glycolytic pyruvate production, redox-dependent lactate dehydrogenase flux and mitochondrial pyruvate metabolism (i.e. PC vs PDH). Lactate concentrations were not significantly different between WT and OB-CON mice but were reduced in fasted mice following pioglitazone treatment (Figure 5B), suggesting that flux from pyruvate into lactate could not explain the decreased pyruvate availability in OB-PIO livers.

**Figure 5:**
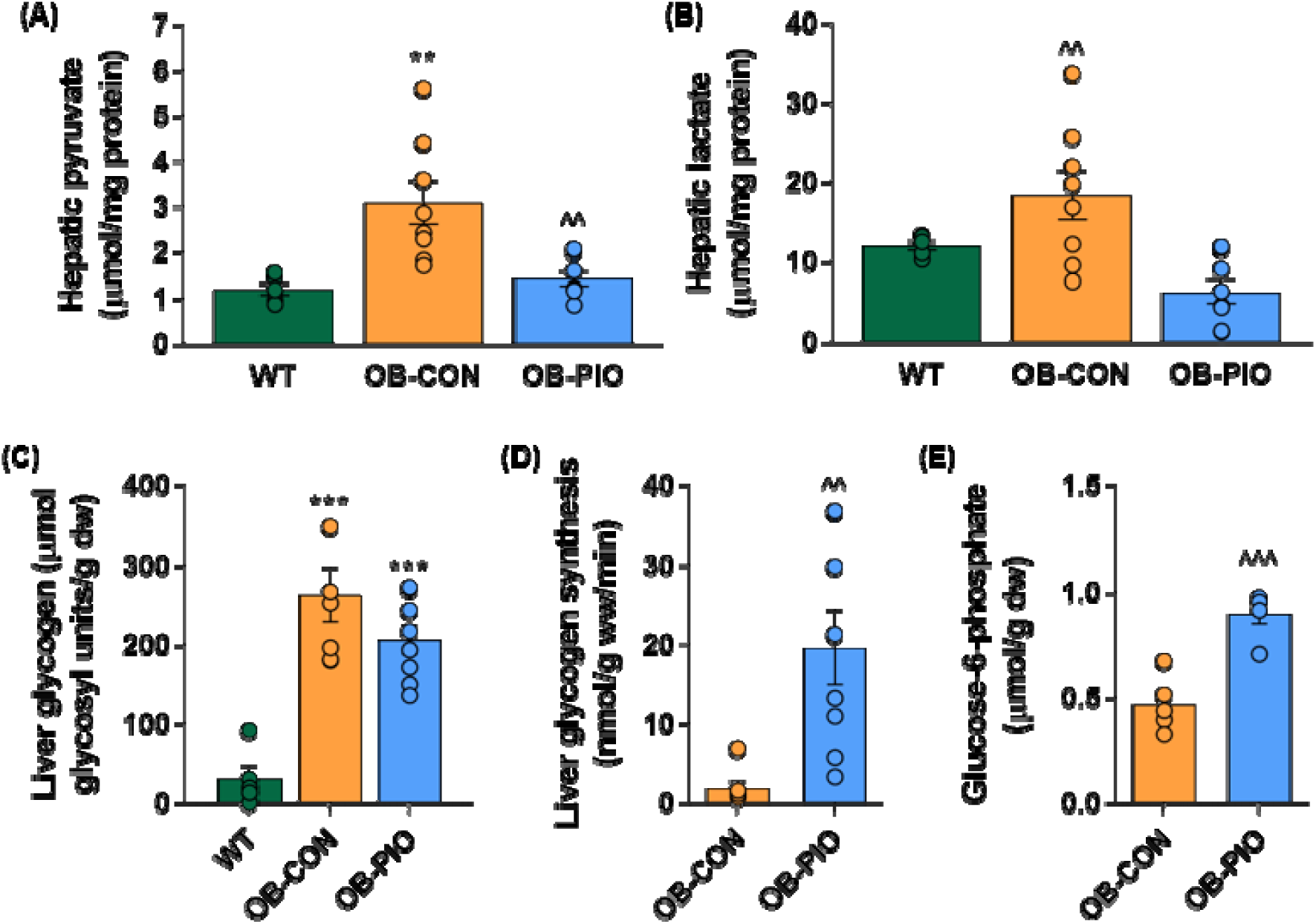
Insulin sensitization is associated with a reduction in glycogenolysis and hepatic pyruvate availability: (A) Hepatic pyruvate, (B) lactate and (C) glycogen concentrations overnight-fasted wild-type (WT), untreated obese (OB-CON) and pioglitazone-treated obese (OB-PIO) mice. (D) Liver glycogen synthesis determined from 3-3H plasma glucose incorporation into hepatic glycogen during insulin clamp in OB-CON and OB-PIO mice. (E) Liver glucose-6-phosphate concentrations in in overnight-fasted OB-CON and OB-PIO mice. **P<0.01, ***P<0.001 vs WT; ^^P<0.01, ^^^P<0.001 vs OB-CON.

Both pyruvate and lactate availability are influenced by liver glycogen metabolism, which is often perturbed in models of obesity and liver disease (32, 33). Indeed, whilst liver glycogen was depleted in fasted WT mice, substantial hepatic glycogen remained in fasted OB-CON and OB-PIO mice (Figure 5C). As such, glycogenolysis accounted for approximately 50% of total glucose production in fasting OB-CON and OB-PIO mice (Figure 2E). Although fasting liver glycogen was no different between OB-CON and OB-PIO mice, scaling the ^2^H NMR date from Figure 2E to absolute rates of glucose production indicated that absolute glycogenolysis was 50% lower in OB-PIO mice compared to OB-CON (0.92 vs 0.60 µmol/min). We subsequently monitored the incorporation of plasma glucose into hepatic glycogen by infusing 3-^3^H glucose under insulin clamp conditions in OB-CON and OB-PIO mice. Consistent with the marked hepatic insulin resistance observed in OB-CON (Supplemental table 1), liver glycogen synthesis from plasma glucose was negligible in these animals (Figure 5D). Insulin sensitization with pioglitazone resulted in a 10-fold increase in the incorporation of plasma glucose into liver glycogen (Figure 5D). Hepatic glucose-6-phosphate concentration, the primary allosteric effector of glycogen metabolism, was also 92% higher in PIO-OB mice (Figure 5E). Thus, excessive glycogen turnover may contribute to the elevation of hepatic pyruvate concentrations in livers from insulin resistant mice.

### Hepatic insulin resistance involves pathological remodeling of mitochondrial lipid species

Improvements in insulin sensitivity (12, 34) and mitochondrial function (19) following prolonged pioglitazone treatment are believed to occur secondarily to reductions in liver fat content. Interestingly however, we found that total hepatic levels of triglyceride were unaltered by four weeks of pioglitazone treatment (Figure 6A). Moreover, related lipid classes such as diacylglycerides (Figure 6B), free fatty acids (Figure 6C) and long-chain acyl-CoAs (Figure 6D) were unaffected or even tended to increase following pioglitazone treatment. Thus, under these conditions, improvements in hepatic insulin sensitivity and glucose homeostasis were dissociated from bulk changes in liver fat content. We subsequently used shotgun lipidomics to investigate additional lipid classes that may be more relevant to the relationship between mitochondrial remodeling and hepatic insulin resistance. Long-chain acylcarnitines (Figure 6E and Supplemental Figure 3A) and ceramides (Figure 6F and Supplemental Figure 3B) are two lipid classes often associated with mitochondrial dysfunction (7, 35) and were both increased in livers from obese mice but were completed normalized by insulin sensitization with pioglitazone. Interestingly, changes in hepatic ceramides were largely driven by N16:0 ceramide (Supplemental Figure 3B), a species that has previously been linked to hepatic insulin resistance, steatosis and mitochondrial impairment (7).

**Figure 6:**
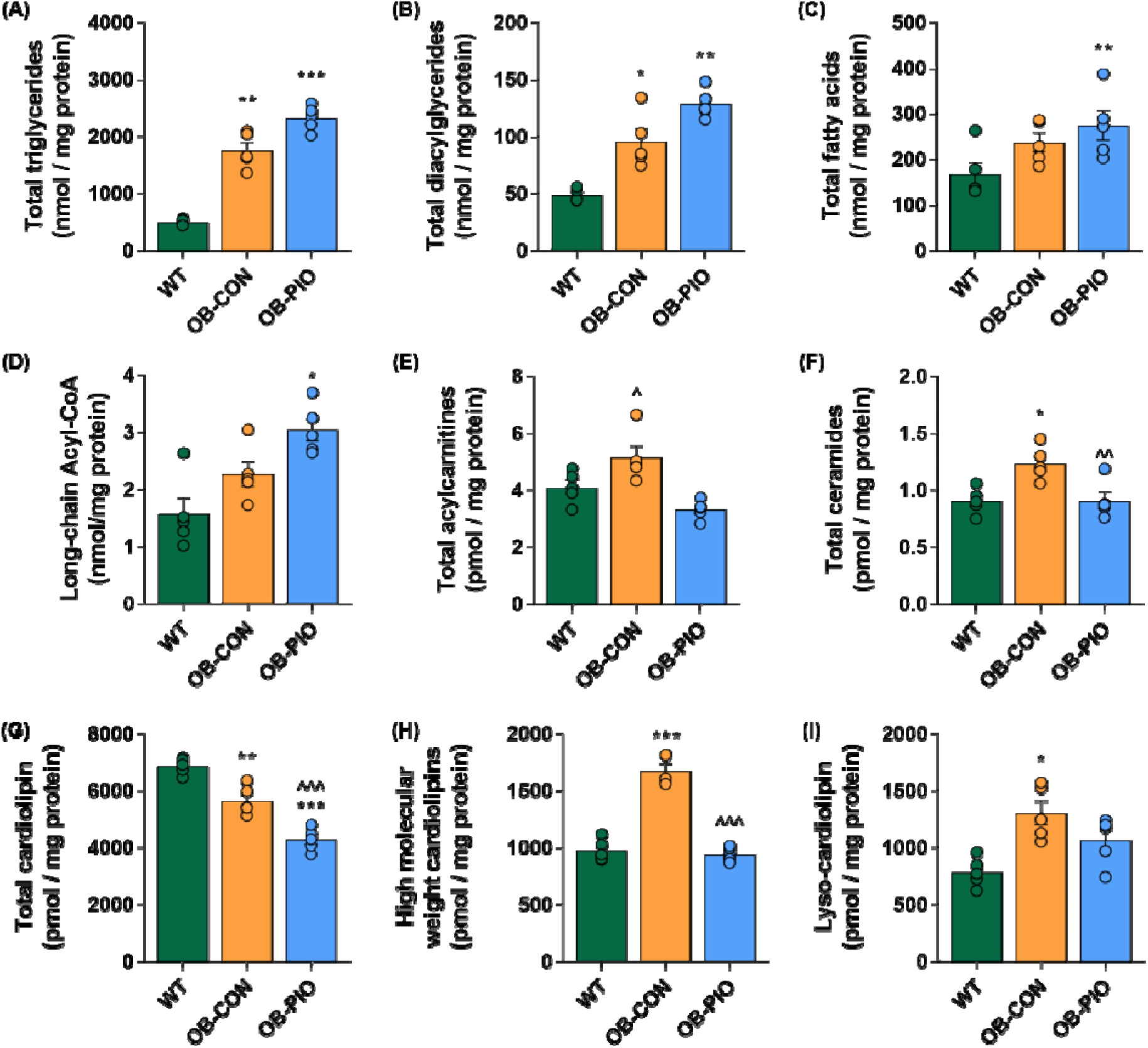
Hepatic insulin resistance involves pathological remodeling of mitochondrial lipid species: Total liver concentrations of (A) triglycerides, (B) diacylglycerides, (C) fatty acids, (D) long-chain acyl-CoAs, (E) long-chain acylcarnitines, (F) ceramides, (G-H) cardiolipins and (I) lyso-cardiolipins calculated by summing individual species determined by mass spectrometry-based shotgun lipidomics, in wild-type, untreated obese mice (OB-CON) and obese mice treated with pioglitazone (∼20 mg/kg/day) for four weeks (OB-PIO). High molecular weight cardiolipins in (H) include cardiolipin species containing fatty acyl chains >C18 (see supplemental Figure 3C). Data are mean ± SE for five mice per group. *P<0.05, **P<0.01, ***P<0.001 vs WT; ^P<0.05, ^^P<0.01, ^^^P<0.001 vs OB-CON.

Another class of lipids thought to play a key role in mitochondrial function are the cardiolipins (36). Total cardiolipin levels were reduced in OB-CON compared to WT mice (Figure 6G), although this difference was driven primarily by fully matured cardiolipin species, including the highly abundant tetralinoleoyl-cardiolipin (Supplemental Figure 3C). Immature cardiolipin species containing 16:0, 16:1 and 18:0 fatty acyl chains, which are related to *de novo* cardiolipin synthesis, were no different between OB-CON and wild-type mice (Supplemental Figure 3C). However, cardiolipin species containing longer fatty acyl chains were dramatically increased in OB-CON mice (Figure 6H and Supplemental Figure 3C). Interestingly, these higher molecular weight cardiolipin species have previously been linked with mitochondrial remodeling and metabolic dysfunction in cardiac tissue (36). Similarly, higher molecular weight lyso-cardiolipins, which act as precursors for cardiolipin synthesis, were also elevated in OB-CON mice compared to WT (Figure 6H and Supplemental Figure 3D). Notably, insulin sensitization partially or completely restored the levels of these higher molecular weight cardiolipin and lyso-cardiolipin species, suggesting a mechanistic role for hepatic cardiolipin remodeling in the pathology of insulin resistance.

## DISCUSSION

Hallmark pathologies of type 2 diabetes and non-alcoholic fatty liver disease, such as excessive glucose output and hepatic steatosis, are underpinned by changes in hepatic mitochondrial function. Here we show that hepatic pyruvate dehydrogenase (PDH) activity is markedly increased in genetically-obese, insulin resistant mice compared to age-matched lean controls. Activation of PDH in livers from obese mice was attributed, in part, to the coordinated suppression of PDK4 and induction of PDP2. Moreover, increases in hepatic TCA cycle activity, which lower acetyl-CoA concentrations, and excessive glycogenolytic flux, which increase pyruvate concentrations, favor a metabolic milieu that promotes allosteric stimulation of hepatic PDH under insulin resistant conditions. These changes were largely reversed by short-term (4 weeks) treatment of obese mice with the insulin sensitizer pioglitazone. Remarkably, these improvements in mitochondrial metabolism and insulin sensitivity occurred without reduction in any of the major hepatic lipid classes. Finally, we provide evidence of a mechanistic role for phospholipid cardiolipin in the link between mitochondrial remodeling and hepatic insulin resistance.

It has been suggested that the accelerated TCA cycle flux in insulin resistant livers reflects the increased energy demands and/or reduced mitochondrial efficiency associated with liver damage (9), although the exact mechanism linking these events has not been delineated. Our data confirm previous suggestions that elevated rates of TCA cycle turnover in NAFLD and NASH are associated with lower concentrations of hepatic acetyl-CoA (9). Consistent with the allosteric regulation of PDH by acetyl-CoA (28), we also show that the resultant reduction in the hepatic acetyl-CoA / CoASH ratio contributes to increased hepatic PDH activation, supporting the increased oxidative demand in these livers. It should be acknowledged that under normal conditions, the contribution of PDH to the hepatic acetyl-CoA pool is relatively minor compared to β-oxidation, such that PDH flux accounts for less than 5% of TCA cycle turnover (37). Therefore, whilst absolute rates of PDH flux may be increased in insulin resistant livers, it is unlikely that this can explain the increased rate of TCA cycle turnover observed in NAFLD and NASH patients. Nevertheless, the accelerated mitochondrial acetyl-group delivery from hyperactive PDH represents an additional metabolic stress on a system that is already overloaded and its suppression in OB-PIO mice is further evidence of the normalized regulation of mitochondrial function with pioglitazone. Interestingly, PDH activity was recently found to be elevated in orthotopic rat hepatomas compared to neighboring liver tissue (38). As such, our data argue that the induction of PDH in insulin resistant livers could represent an important, targetable event in the progression from NAFLD and NASH to hepatocellular carcinoma.

In addition to changes in acetyl-CoA concentrations, the increased PDH activation in obese livers was driven by an increase in the PDH E1α subunit, as well as reduced PDK4 and increased PDP2 protein levels, favoring covalent activation of this subunit. Pioglitazone restored protein levels of PDK4 and PDP2 independent of total E1α protein expression. PDK4 is induced by physiological states of low insulin, including starvation and type I diabetes (39, 40), whereas PDP2 is suppressed under such conditions (27). Therefore, the reciprocal changes in PDP2 observed in OB-CON and OB-PIO mice may be explained by the pioglitazone-mediated improvement in peripheral insulin sensitivity, resulting in lower circulating insulin concentrations. However, PDK4 was also induced by pioglitazone in isolated hepatocytes, suggesting a more direct mode of action. Interestingly, RNA-seq analysis of pioglitazone-treated hepatocytes revealed PDK4 to be the most highly-upregulated gene (unpublished data). The direct induction of PDK4 by pioglitazone is consistent with previous observations that pioglitazone acutely inhibits pyruvate oxidation (16, 41). Although pioglitazone had no effect on PDH activation in perfused livers, these experiments were conducted using 1 mM pyruvate to optimize glucose production. High pyruvate concentrations are known to stimulate PDH flux by inhibiting PDH kinases (31) and may have subsequently masked the effects of pioglitazone on PDH activation under these conditions.

Our findings also suggest that the accelerated rates of oxidative metabolism seen in insulin resistant livers are supported by changes in mitochondrial number, structure and function. Intuitively, mitochondrial substrate delivery or partitioning must also be adapted to meet the increased demands of greater TCA cycle turnover. It has been argued that the primary source of this increased substrate delivery is from adipose tissue lipolysis (42) and indeed, the accumulation of long-chain acylcarnitine species observed in the livers of obese mice here indicates excessive mitochondrial lipid delivery. However, our data also suggests that the effects of pioglitazone on mitochondrial function may be distinct from actions on adipose tissue. This is supported by the lack of change in circulating free fatty acids and glycerol, suggesting unaltered lipolysis, as well as the increases in hepatic triglyceride and diacylglyceride contents in OB-PIO mice compared to OB-CON. Instead, we show that disturbed hepatic glycogen metabolism plays an important role in the oversupply of mitochondrial substrates. Insulin resistant livers were characterized by excessive glycogen storage, persistent glycogenolysis and impaired glycogen synthesis in response to insulin infusion. Pioglitazone restored insulin-stimulated glycogen synthesis and suppressed glycogenolytic flux, which assisted the lowering of hepatic glucose production but also likely restricted glycolytic pyruvate generation. Our observation that hepatic pyruvate concentration was elevated in obese mice is consistent with the finding that circulating pyruvate concentrations are increased in NAFLD patients (9). Reducing hepatic pyruvate availability may thus represent a central node linking insulin sensitization with the improvement of mitochondrial function, by relieving stimulation of both PDH and PC-mediated TCA cycle substrate delivery.

A previous study found that chronic (6 months) pioglitazone treatment suppressed hepatic TCA cycle flux in a rodent model of NASH, attributing the effects to reduced liver accumulation of triglycerides and diacylglycerides (19). However, no detailed mechanism was identified for the alteration of mitochondrial function by pioglitazone. Here we confirm the ability of pioglitazone to slow TCA cycle activity by ^13^C NMR and corroborate this with pioglitazone-induced decreases in citrate synthase activity, protein and lipid markers of mitochondrial content and function. However, we demonstrate that this occurs rapidly (within four weeks of treatment) and either precedes or is dissociated from any effects on liver fat content, as major classes of hepatic lipids (triglycerides, diacylglycerides and free fatty acids) were unchanged by pioglitazone treatment. Instead, we present evidence that the rescue of mitochondrial function by pioglitazone involves remodeling of the mitochondrial lipid cardiolipin. Cardiolipins are a class of inner mitochondrial membrane phospholipids which function to maintain cellular bioenergetics by regulating the organization of mitochondrial proteins (36). Human data suggest that hepatic levels of total cardiolipin are unchanged (43) or modestly increased (7) in NAFLD and NASH patients. However, the fatty acyl-chain composition of cardiolipin appears to be more important than total content in determining its impact upon hepatic mitochondrial function (44). As such, a major strength of this work is the use of high-resolution shotgun lipidomics to identify individual cardiolipin species. Our approach revealed a marked reduction in the most abundant cardiolipin species in livers from obese mice, but an increase in many of the less abundant species, particularly those of longer acyl chain length. This pattern of hepatic cardiolipin remodeling recapitulates findings from insulin resistant cardiac tissue (45) and is reported to promote oxidative stress and mitochondrial dysfunction by rendering cardiolipin more susceptible to peroxidation (46). Therefore, it seems likely that changes in the content and acyl-chain composition of hepatic cardiolipin contribute to the excessive TCA cycle activity and oxidative burden observed in insulin resistant livers. As such, reducing longer acyl-chain cardiolipin species may represent an important mechanism through which insulin sensitization with pioglitazone can protect mitochondria from lipid peroxidation and thus preserve hepatic mitochondrial function.

Two additional classes of lipids that have been closely linked with mitochondrial function in insulin resistant tissues are long-chain acylcarnitines and ceramides. Acylcarnitines are synthesized by carnitine palmitoyltransferase I (CPTI) to facilitate transport of fatty acids into the mitochondria for subsequent β-oxidation (47). In agreement with evidence from human NASH patients (7, 48), total acylcarnitine content was elevated in OB-CON livers, reflecting an imbalance between CPTI-mediated acylcarnitine production and their disposal via β-oxidation. Rates of complete β-oxidation (acetyl-CoA production) were recently shown to be normal in patients with varying severities of NAFLD (9) and therefore it seems likely that the accumulation of hepatic long-chain acylcarnitines reflects increased mitochondrial fatty acid delivery, rather than impaired oxidation *per se*. Interestingly, the pioglitazone-induced reduction in acylcarnitines was mirrored by a reduction in hepatic ceramide concentrations, a finding that is consistent with a recent study which reported a lowering of plasma ceramide concentrations in pioglitazone-treated human subjects with metabolic syndrome (49). Since the patterns of ceramide and cardiolipin remodeling observed here have been directly linked with mitochondrial dysfunction and impaired hepatic β-oxidation (35) (45), it is tempting to speculate that the resolution of hepatic acylcarnitine concentrations reflects ceramide and/or cardiolipin-mediated improvements in mitochondrial function. It is also particularly noteworthy that these beneficial alterations in less-abundant hepatic lipid species occurred despite increases in liver triglycerides and diacylglycerides, highlighting the ability of pioglitazone to protect mitochondria from lipid overload even in the setting of increased hepatic lipid storage.

In conclusion, we identify pyruvate dehydrogenase activation and cardiolipin remodeling as novel features of hepatic mitochondrial dysfunction in insulin resistant obese mice. Treatment with pioglitazone reversed these maladaptive processes and improved whole-body glucose metabolism, without reducing hepatic triglyceride levels, suggesting that the link between hepatic lipid storage, mitochondrial function and insulin sensitivity is more nuanced than previously assumed. Hepatic Insulin sensitization drives multiple pathways to relieve the oxidative burden of excessive TCA cycle flux and improve mitochondrial function, highlighting the need to target insulin resistance as a primary treatment for liver disease.

## ACKNOWLEDGEMENTS

Project supported by funding from the University of Florida’s Southeast Center for Integrated Metabolomics through grant number U24DK097209 from the National Institute of Health’s Common Fund metabolomics program. A portion of this work was performed at the National High Magnetic Field Laboratory, which is supported by National Science Foundation Cooperative Agreement number DMR-1644779, & the State of Florida, NIH P41-122698, and NIH R01-105346. C.E.S. is supported by an UT Health San Antonio OPA Postdoctoral Research Fellowship

## AUTHOR CONTRIBUTIONS

Conceptualization, C.E.S., M.R., M.M., and L.N.; Methodology, C.E.S., M.R., M.M., J.P.P., E.S.J., C.R.M., X.H., and L.N.; Formal analysis, C.E.S., M.R., and J.P.P.; Investigation, C.E.S., M.R., M.F., and T.B.; Writing – Original Draft, C.E.S., and L.N.; Writing – Review and Editing, C.E.S., M.R., M.M., J.P.P., M.M., and L.N.; Supervision, M.M., X.H., and L.N.; Funding Acquisition, L.N.

## SUPPLEMENTAL INFORMATION

**Supplemental Table 1:**
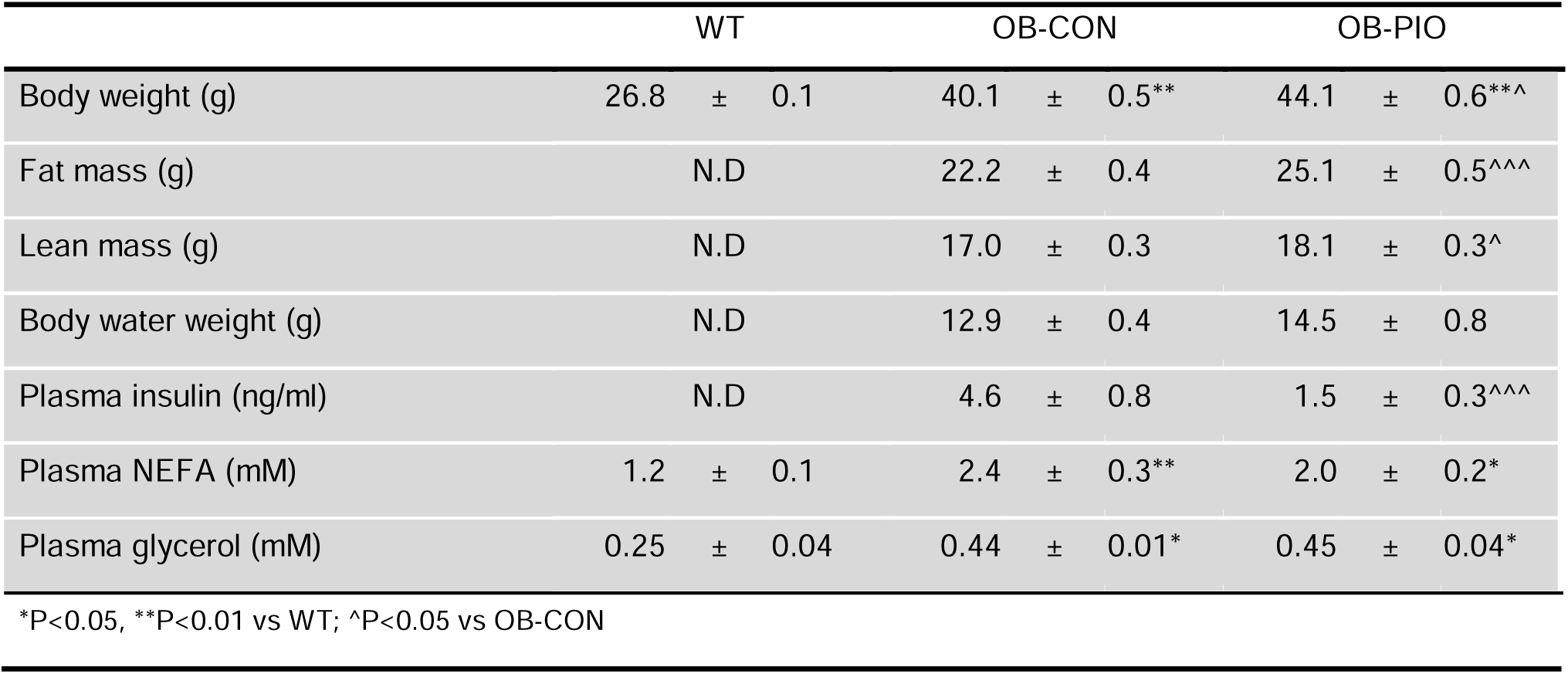
Metabolic parameters of wild-type (WT) and Ob/Ob mice treated with placebo (OB-CON) or 25 mg/kg/day pioglitazone (OB-PIO) for 4 weeks

## SUPPLEMENTAL FIGURES

**Supplemental Figure 1:**
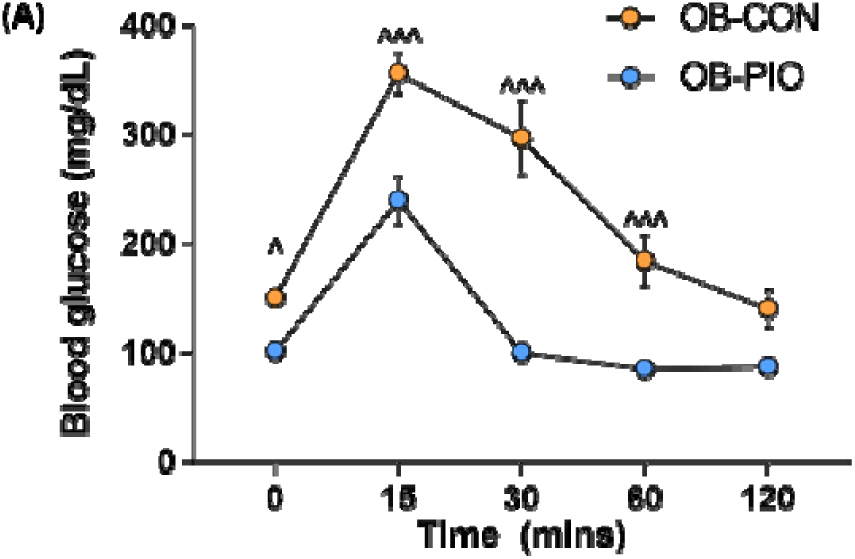
(A) Blood glucose during oral glucose tolerance test in untreated obese mice (OB-CON) and obese mice treated with pioglitazone (∼20 mg/kg/day) for two weeks (OB-PIO).

**Supplemental Figure 2:**
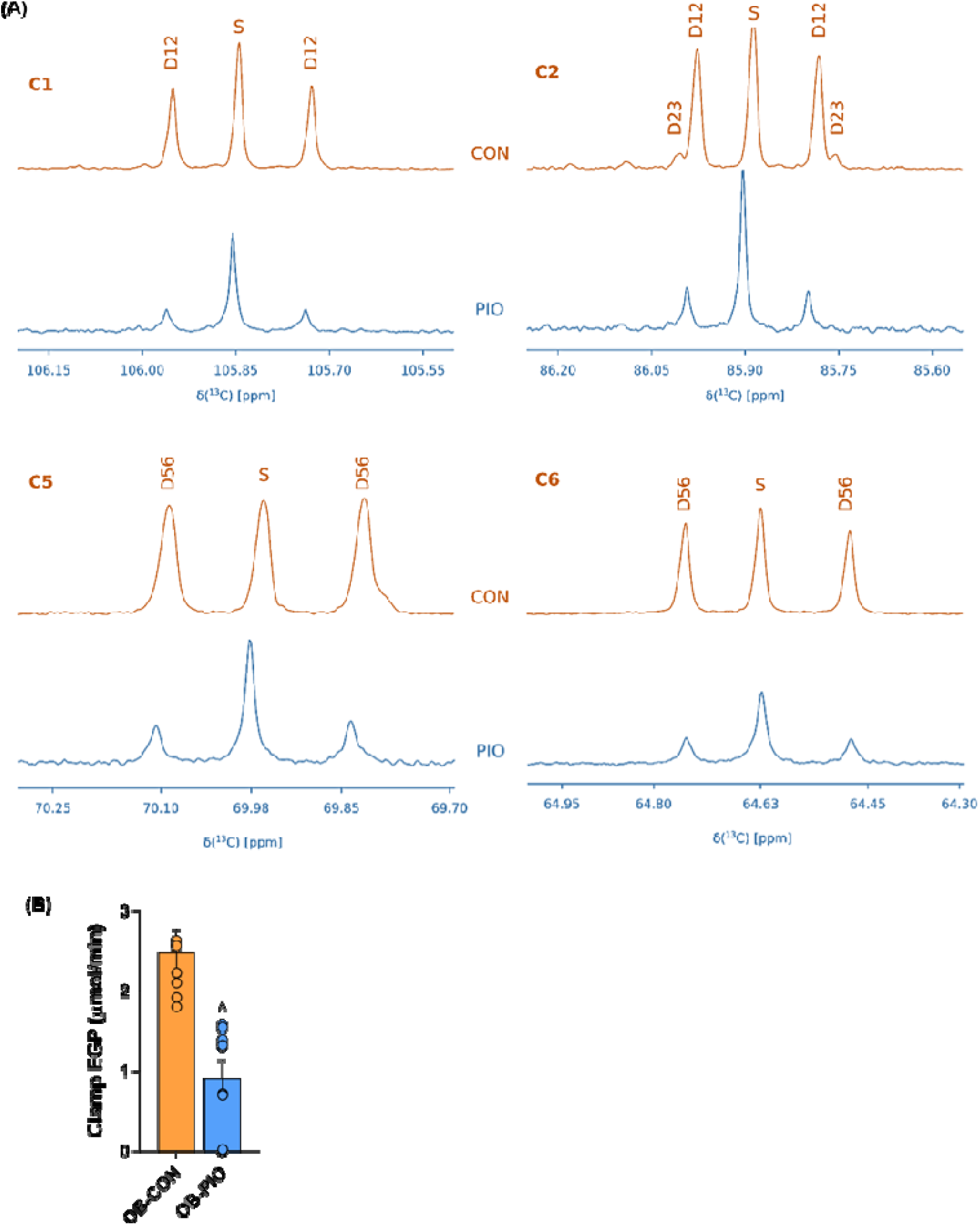
(A) ^13^C spectra showing C-1, C-2, C-5 and C-6 resonance of monoacetone glucose synthesized from plasma in OB-CON (orange) compared to OB-PIO (blue) mice. (B) Endogenous glucose production in overnight-fasted OB-CON (n=7) and OB-PIO (n=8) mice measured using 3-^3^H glucose. ^P<0.05 vs OB-CON.

**Supplemental Figure 3:**
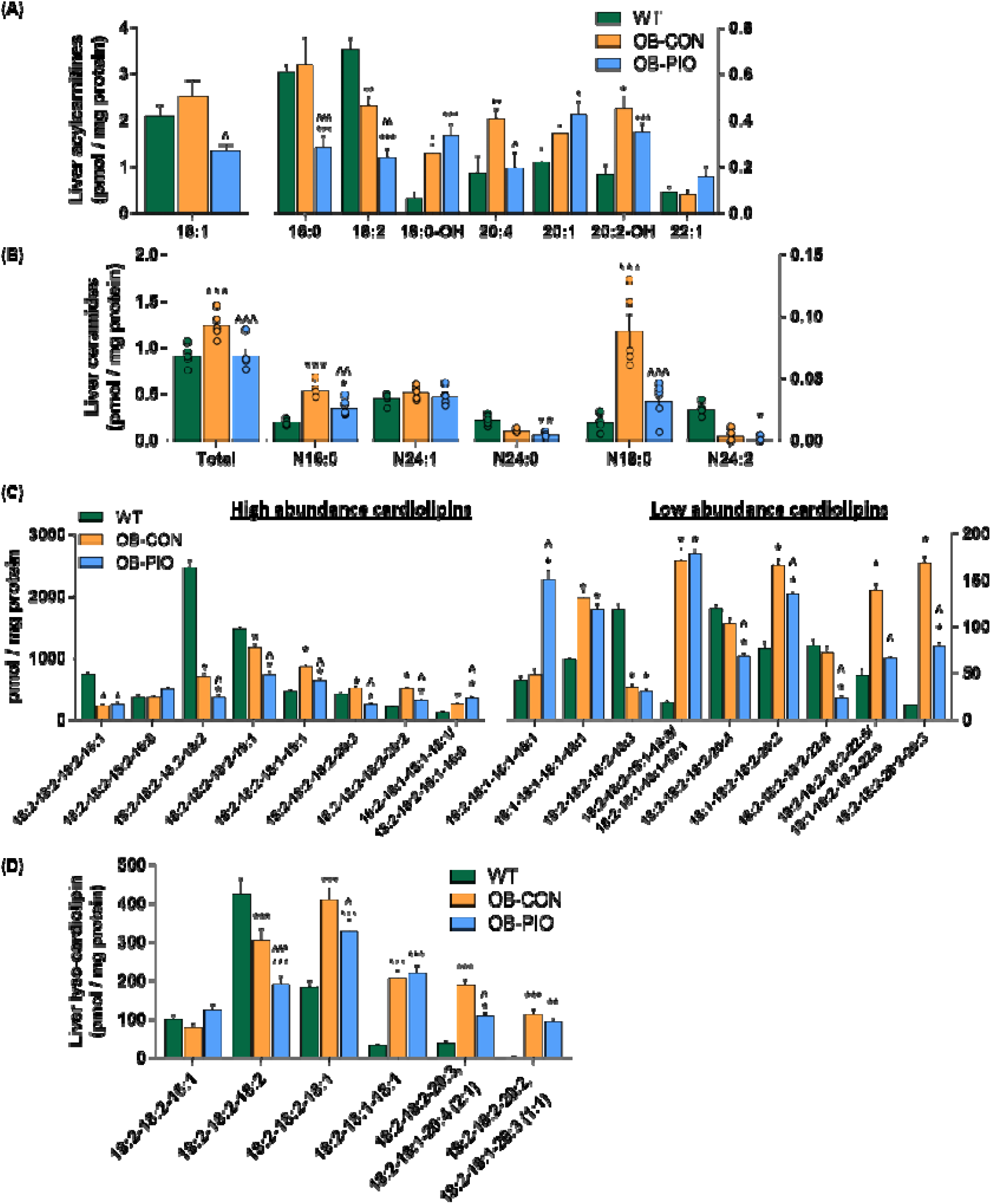
Hepatic concentrations of individual species of (A) long-chain acylcarnitines, (B) ceramides, (C) cardiolipins and (D) lyso-cardiolipins in untreated wild-type, OB-CON and OB-PIO mice, measured by mass spectrometry-based shotgun lipidomics. *P<0.05, **P<0.01, ***P<0.001 vs WT; ^P<0.05, ^^^P<0.001 vs OB-CON.

## METHODS

### Experimental Model and Subject Details

#### Animal Studies

Eight-week old wild type (C57BL/6J; JAX #000664) and Ob/Ob (B6 ob; JAX #000632) male mice were purchased from The Jackson Laboratory, housed in environmentally-controlled conditions (23°C, 12 hour light/dark cycles) and provided ad-libitum access to water and a control (70% kcal carbohydrate, 10% kcal fat, 20% kcal protein; D12450J; Research Diets Inc.) or macronutrient-matched diet supplemented with pioglitazone (300 mg/kg; D17010703; Research Diets Inc.). Experiments were conducted in 12-14-week aged mice following four weeks of diet treatments. All procedures were approved by the Institutional Animal Care and Use Committee at University of Texas Health Science Center at San Antonio.

#### Cell Culture

Primary hepatocytes were isolated from 12-16-week old wild type male mice and maintained in serum-free DMEM supplemented with Pen-Strep (50 U/ml; Gibco). 5×10^5^ cells were seeded in collagen-coated 6-well plates (354400; Corning) and treated with pioglitazone (2.5 – 10µM) or vehicle control (DMSO) for 24-48 hours. Treatments were added 24 hours after plating.

## Method Details

### Oral Glucose Tolerance Test

Overnight-fasted (16 hours) Ob/Ob mice fed control (OB-CON) or pioglitazone (OB-PIO) diet were administered with a fixed dose of 50 mg glucose (250µl D20 20% dextrose) by oral gavage. Glucose concentration was determined in whole-blood sampled at baseline and at 15-, 30-, 60- and 120-minute intervals from a tail incision using a glucometer (Contour next EZ).

### Endogenous Glucose Production

Overnight-fasted OB-CON and OB-PIO mice were infused with 3-^3^H glucose (0.1 µCi / min) for two hours through a jugular vein catheter that had been inserted five days prior to experiments. Blood was sampled at −20, −10, 0, 90, 100, 110 and 120 minutes for the measurement of plasma glucose specific activity.

### Peripheral Insulin Sensitivity

For euglycemic-hyperinsulinemic clamp studies in OB-CON and OB-PIO mice, insulin was infused in a primed-continuous manner (50 mU/kg bolus then 5 mU/kg/min) and blood was sampled every five minutes to monitor glucose concentrations. Glucose (20% glucose) was infused at a variable rate to maintain euglycemia at 100 mg/dL and red blood cells from donor mice were infused continuously to prevent anemic volume depletion. A 3-^3^H glucose tracer was infused continuously (0.1 µCi / min) for the determination of insulin-stimulated glucose output.

### Body Composition

Lean mass, fat mass and total body water were determined in OB-CON and OB-PIO mice by qMRI at the San Antonio Nathan Shock Centre Aging Animal Models and Longevity Assessment Core.

### Plasma Analytes

Plasma insulin was determined by ELISA (#90080 Crystal Chem). Plasma free fatty acids (999-34691 Wako), glycerol (F6428 Sigma) and triglycerides (TR22421 ThermoFisher) were determined by enzymatic calorimetric assay.

### Liver Citrate Synthase Activity

Citrate synthase activity was determined in liver homogenates by spectrophotometric assay which monitors the rate of CoASH formation from the condensation of oxaloacetate with acetyl-CoA in the presence of the 5,5-dithio-bis-2-nitrobenzoic acid (50). Data were normalized to homogenate protein content (BCA assay; ThermoFisher).

### Western Blotting and Quantitative Real-Time PCR (qRT-PCR)

Immunoblot analysis was carried out on cell or liver lysates using primary antibodies from Cell Signaling Technologies and Abcam (described in Key Resources Table) and developed by chemiluminescence method (ECL). Quantitative real-time PCR (qRTPCR) was performed using pre-designed TaqMan probes (Life Technologies). Data were normalized to reference genes B2m and HMBS.

### Multi-dimensional Mass Spectrometry-based Shotgun Lipidomics

Liver tissue was lyophilized and homogenized in ice-cold diluted phosphate-buffered saline (0.1X PBS). Lipids were extracted by a modified procedure of Bligh and Dyer extraction in the presence of internal standards, which were added based on total protein content as previously described (51, 52). A triple-quadrupole mass spectrometer (Thermo Scientific TSQ Altis, CA, USA) and a Quadrupole-Orbitrap™ mass spectrometer (Thermo Q Exactive™, San Jose, CA) equipped with a Nanomate device (Advion Bioscience Ltd., NY, USA) and Xcalibur system software was used as previously described (53). Briefly, diluted lipid extracts were directly infused into the electrospray ionization source through a Nanomate device. Signals were averaged over a 1-min period in the profile mode for each full scan MS spectrum. For tandem MS, a collision gas pressure was set at 1.0 mTorr, but the collision energy varied with the classes of lipids. Similarly, a 2-to 5-min period of signal averaging in the profile mode was employed for each tandem MS mass spectrum. All full and tandem MS mass spectra were automatically acquired using a customized sequence subroutine operated under Xcalibur software. Data processing including ion peak selection, baseline correction, data transfer, peak intensity comparison, 13C deisotoping, and quantitation were conducted using a custom programmed Microsoft Excel macro as previously described after considering the principles of lipidomics (54).

### ^13^C and ^2^H tracer studies

For the assessment of fractional glucose production and mitochondrial pyruvate metabolism, overnight-fasted mice were administered with 27 µl/g body weight ^2^H_2_O (99.9%; Cambridge Isotopes) by intraperitoneal injection. Thirty minutes after the ^2^H_2_O injection, mice were anaesthetized with isoflurane and injected with 30 µmols [2,3-^13^C] pyruvate (Cambridge Isotopes) in 150 µl saline. Eight minutes after [2,3-^13^C] pyruvate injection, livers were freeze-clamped *in situ* and blood was collected by cardiac puncture. A portion of liver was pulverized under liquid nitrogen, extracted in ice-cold perchloric acid, neutralized with potassium hydroxide and lyophilized in preparation for glutamate isotopomer analysis. Plasma was isolated from whole blood by centrifugation and used for the preparation of monoacetone glucose.

### ^13^C NMR Analysis of Liver Glutamate Isotopomers

^13^C NMR spectra were acquired using an NMR spectrometer interfaced with a 14.1 T magnet equipped with a home-built superconducting (HTS) probe (55). NMR samples were prepared by dissolving the lyophilized powder from PCA extraction in 90% (v/v) of 50 mM sodium phosphate in D_2_O (pH 7.0) containing 2 mM EDTA. To this solution, 10% (v/v) internal standard containing 5 mM DSS-d6 and 0.2% (w/v) NaN_3_ in D_2_O was added. ^13^C spectra were measured with a spectra width of 240 ppm and an acquisition time of 1.5 s at 298 K. ^1H^decoupling was performed using WALTZ-16 scheme. ^13^C NMR spectra were processed (zero filled to 131072 points, 0.5 Hz exponential line broadening, and polynomial baseline correction) using Mestrenova (v13.0 or above). Chemical shift referencing was carried out by setting taurine resonance to 48.4 ppm. Glutamate resonances were identified and fitted to mixed Gaussian/Lorentzian line-shapes and peak areas were obtained.

### Synthesis of Monoacetone Glucose

To the plasma, barium hydroxide (0.05 M) and zinc sulphate (0.1 M) solutions were added, vortexed and centrifuged. The supernatant was separated and lyophilized. To the lyophilized powder, a mixture of acetone and concentrated sulfuric acid was added, and vortexed for 4 hours. After 4 hours, 1.5 M sodium carbonate was added to adjust the pH to 1.75 and stirred for 24 hours. This solution is then lyophilized, monoacetone glucose (MAG) is extracted with ethyl acetate and dried under a stream of nitrogen.

### ^13^C and ^2^H NMR Analysis of Monoacetone Glucose

Samples of synthesized MAG were dissolved in a mixture containing 94% (v/v) CD3CN and 6% (v/v) D2O. ^13^C NMR spectra of MAG were recorded and processed with same parameters as those used for recording the PCA extracted samples. Chemical shift referencing was carried out by setting the central acetonitrile resonance to 1.3 ppm. All other processing steps were identical to those used to analyze glutamate isotopomers. ^2^H NMR spectra were measured using an NMR spectrometer (14.1 T magnet) equipped with a 3 mm broadband probe with a spectral width of 10.5 ppm and an acquisition time of 1 s at 323 K. ^2^H NMR spectra were processed using a 2 Hz exponential line broadening and polynomial baseline correction. Chemical shifts were referenced by setting the acetonitrile resonances to 1.97 ppm. Fractional contributions to glucose production by the livers were estimated using a Bayesian approach described elsewhere (56).

### Pyruvate Dehydrogenase Activation Status

The activation status of pyruvate dehydrogenase was assayed in liver homogenate from a portion of the left lobe of the liver that had been freeze-clamped *in situ* in isoflurane-anaesthetized mice. The rate of acetyl-CoA production from pyruvate was monitored over 10 minutes at 37° in a buffer containing NAD^+^, Coenzyme A and thiamine pyrophosphate, as well as the PDK and PDP inhibitors dichloroacetate and sodium fluoride, respectively (57).

### Perfused Liver Studies

Livers were isolated from fed WT mice following ketamine / xylazine anesthesia and perfused through a portal vein catheter using a peristaltic pump without recirculation. HBSS buffer containing 1mM pyruvate / 10 mM lactate and bubbled continuously with 95% oxygen / 5% CO_2_ was perfused at a rate of ∼4 ml/min. After an equilibration period of 10 minutes, vehicle (DMOS) or Pioglitazone (final concentration 10µM) was added to the perfusion media. Effluent was sampled at 10-minute intervals for glucose measurement (A22189 ThermoFisher) and a portion of liver was freeze-clamped in liquid nitrogen-cooled tongs at the end of the perfusion to determine pyruvate dehydrogenase activation.

### Liver Metabolite Analyses

A ∼50 mg portion of freeze-clamped liver from isoflurane-anaesthetized mice was lyophilized overnight metabolites and extracted in 0.5 M perchloric acid followed by alkaline digestion for the determination of acid-soluble metabolites. Acetyl-CoA and CoASH were assayed by radioenzymatic assay (29). Lactate was assayed by the fluorometric detection of NADH in the presence of excess concentrations of NAD^+^, hydrazine and lactate dehydrogenase (58). Glucose-6-phosphate (G6P) was assayed by the fluorometric detection of NADH in the presence of 50 mmol/L triethanolamine, 0.5 mmol/L dithiothreitol, 0.25 mmol/L ATP, 1 mmol/L NAD, and 0.6 units bacterial G6PDH (G5760; Sigma-Aldrich).

### Hepatic Glycogen Synthesis

Livers harvested from anaesthetized mice following 3-^3^H infusion during euglycemic clamp were used to estimate rates of plasma glucose incorporation into liver glycogen. A ∼20 mg portion of liver was alkaline-digested at 85° for 30 minutes, allowed to cool and neutralized with hydrochloric acid. The neutralized extract was split in half and one half treated with amyloglucosidase for one hour at room temperature. The liberated glucose was subsequently measured in an aliquot of each extract. Five volumes of ethanol were added to the remaining extract to precipitate glycogen. The glycogen pellet was further washed in ethanol, solubilized in water and added to scintillation vials for radioactivity counting. The specific activity was measured as DPM / glycogen and was normalized to plasma tritium specific activity to estimate glycogen synthesis rates during the entire clamp period.

## Quantification and Statistical Analyses

Differences between WT, OB-CON and OB-PIO groups were assessed using one-way analysis of variance with individual differences isolated post-hoc using the Tukey test. For analyses where no wild-type group were included, differences between OB-CON and OB-PIO were assessed using unpaired t-testing. Significance testing was performed using GraphPad Prism 7. Data are presented as mean ± standard error and data were considered significantly different at P<0.05.

## REFERENCES

1. Bril F, Cusi K. Management of Nonalcoholic Fatty Liver Disease in Patients With Type 2 Diabetes: A Call to Action. Diabetes Care. 2017;40(3):419–30.

2. Wang C, Wang X, Gong G, Ben Q, Qiu W, Chen Y, et al. Increased risk of hepatocellular carcinoma in patients with diabetes mellitus: a systematic review and meta-analysis of cohort studies. Int J Cancer. 2012;130(7):1639–48.

3. Bedi O, Aggarwal S, Trehanpati N, Ramakrishna G, Krishan P. Molecular and Pathological Events Involved in the Pathogenesis of Diabetes-Associated Nonalcoholic Fatty Liver Disease. J Clin Exp Hepatol. 2019;9(5):607–18.

4. Gaggini M, Morelli M, Buzzigoli E, DeFronzo RA, Bugianesi E, Gastaldelli A. Non-alcoholic fatty liver disease (NAFLD) and its connection with insulin resistance, dyslipidemia, atherosclerosis and coronary heart disease. Nutrients. 2013;5(5):1544–60.

5. Sunny NE, Parks EJ, Browning JD, Burgess SC. Excessive hepatic mitochondrial TCA cycle and gluconeogenesis in humans with nonalcoholic fatty liver disease. Cell Metab. 2011;14(6):804–10.

6. Schmid AI, Szendroedi J, Chmelik M, Krssák M, Moser E, Roden M. Liver ATP synthesis is lower and relates to insulin sensitivity in patients with type 2 diabetes. Diabetes Care. 2011;34(2):448–53.

7. Peng KY, Watt MJ, Rensen S, Greve JW, Huynh K, Jayawardana KS, et al. Mitochondrial dysfunction-related lipid changes occur in nonalcoholic fatty liver disease progression. J Lipid Res. 2018;59(10):1977–86.

8. Koliaki C, Szendroedi J, Kaul K, Jelenik T, Nowotny P, Jankowiak F, et al. Adaptation of hepatic mitochondrial function in humans with non-alcoholic fatty liver is lost in steatohepatitis. Cell Metab. 2015;21(5):739–46.

9. Fletcher JA, Deja S, Satapati S, Fu X, Burgess SC, Browning JD. Impaired ketogenesis and increased acetyl-CoA oxidation promote hyperglycemia in human fatty liver. JCI Insight. 2019;5.

10. Miyazaki Y, Mahankali A, Matsuda M, Glass L, Mahankali S, Ferrannini E, et al. Improved glycemic control and enhanced insulin sensitivity in type 2 diabetic subjects treated with pioglitazone. Diabetes Care. 2001;24(4):710–9.

11. Bril F, Kalavalapalli S, Clark VC, Lomonaco R, Soldevila-Pico C, Liu IC, et al. Response to Pioglitazone in Patients With Nonalcoholic Steatohepatitis With vs Without Type 2 Diabetes. Clin Gastroenterol Hepatol. 2018;16(4):558–66 e2.

12. Cusi K, Orsak B, Bril F, Lomonaco R, Hecht J, Ortiz-Lopez C, et al. Long-Term Pioglitazone Treatment for Patients With Nonalcoholic Steatohepatitis and Prediabetes or Type 2 Diabetes Mellitus: A Randomized Trial. Ann Intern Med. 2016;165(5):305–15.

13. Spiegelman BM. PPAR-gamma: adipogenic regulator and thiazolidinedione receptor. Diabetes. 1998;47(4):507–14.

14. Bays H, Mandarino L, DeFronzo RA. Role of the adipocyte, free fatty acids, and ectopic fat in pathogenesis of type 2 diabetes mellitus: peroxisomal proliferator-activated receptor agonists provide a rational therapeutic approach. J Clin Endocrinol Metab. 2004;89(2):463–78.

15. Miyazaki Y, DeFronzo RA. Rosiglitazone and pioglitazone similarly improve insulin sensitivity and secretion, glucose tolerance and adipocytokines in type 2 diabetic patients. Diabetes Obes Metab. 2008;10(12):1204–11.

16. Shannon CE, Daniele G, Galindo C, Abdul-Ghani MA, DeFronzo RA, Norton L. Pioglitazone inhibits mitochondrial pyruvate metabolism and glucose production in hepatocytes. FEBS J. 2017;284(3):451–65.

17. Raman P, Judd RL. Role of glucose and insulin in thiazolidinedione-induced alterations in hepatic gluconeogenesis. Eur J Pharmacol. 2000;409(1):19–29.

18. Nishimura Y, Inoue Y, Takeuchi H, Oka Y. Acute effects of pioglitazone on glucose metabolism in perfused rat liver. Acta Diabetol. 1997;34(3):206–10.

19. Kalavalapalli S, Bril F, Koelmel JP, Abdo K, Guingab J, Andrews P, et al. Pioglitazone improves hepatic mitochondrial function in a mouse model of nonalcoholic steatohepatitis. Am J Physiol Endocrinol Metab. 2018;315(2):E163–E73.

20. McCommis KS, Chen Z, Fu X, McDonald WG, Colca JR, Kletzien RF, et al. Loss of Mitochondrial Pyruvate Carrier 2 in the Liver Leads to Defects in Gluconeogenesis and Compensation via Pyruvate-Alanine Cycling. Cell Metab. 2015;22(4):682–94.

21. Garcia J, Decker CW, Sanchez SJ, Ouk JM, Siu KM, Han D. Obesity and steatosis promotes mitochondrial remodeling that enhances respiratory capacity in the liver of ob/ob mice. FEBS Lett. 2018;592(6):916–27.

22. Aronoff S, Rosenblatt S, Braithwaite S, Egan JW, Mathisen AL, Schneider RL. Pioglitazone hydrochloride monotherapy improves glycemic control in the treatment of patients with type 2 diabetes: a 6-month randomized placebo-controlled dose-response study. The Pioglitazone 001 Study Group. Diabetes Care. 2000;23(11):1605–11.

23. Jin ES, Moreno KX, Wang JX, Fidelino L, Merritt ME, Sherry AD, et al. Metabolism of hyperpolarized [1-(13)C]pyruvate through alternate pathways in rat liver. NMR Biomed. 2016;29(4):466–74.

24. Sherry AD, Jeffrey FM, Malloy CR. Analytical solutions for (13)C isotopomer analysis of complex metabolic conditions: substrate oxidation, multiple pyruvate cycles, and gluconeogenesis. Metab Eng. 2004;6(1):12–24.

25. Jeoung NH, Wu P, Joshi MA, Jaskiewicz J, Bock CB, Depaoli-Roach AA, et al. Role of pyruvate dehydrogenase kinase isoenzyme 4 (PDHK4) in glucose homoeostasis during starvation. Biochem J. 2006;397(3):417–25.

26. Huang B, Gudi R, Wu P, Harris RA, Hamilton J, Popov KM. Isoenzymes of pyruvate dehydrogenase phosphatase. DNA-derived amino acid sequences, expression, and regulation. J Biol Chem. 1998;273(28):17680–8.

27. Huang B, Wu P, Popov KM, Harris RA. Starvation and diabetes reduce the amount of pyruvate dehydrogenase phosphatase in rat heart and kidney. Diabetes. 2003;52(6):1371–6.

28. Batenburg JJ, Olson MS. Regulation of pyruvate dehydrogenase by fatty acid in isolated rat liver mitochondria. J Biol Chem. 1976;251(5):1364–70.

29. Cederblad G, Carlin JI, Constantin-Teodosiu D, Harper P, Hultman E. Radioisotopic assays of CoASH and carnitine and their acetylated forms in human skeletal muscle. Anal Biochem. 1990;185(2):274–8.

30. Carlin JI, Harris RC, Cederblad G, Constantin-Teodosiu D, Snow DH, Hultman E. Association between muscle acetyl-CoA and acetylcarnitine levels in the exercising horse. J Appl Physiol (1985). 1990;69(1):42–5.

31. Cate RL, Roche TE. A unifying mechanism for stimulation of mammalian pyruvate dehydrogenase(a) kinase by reduced nicotinamide adenine dinucleotide, dihydrolipoamide, acetyl coenzyme A, or pyruvate. J Biol Chem. 1978;253(2):496–503.

32. d’Avignon DA, Puchalska P, Ercal B, Chang Y, Martin SE, Graham MJ, et al. Hepatic ketogenic insufficiency reprograms hepatic glycogen metabolism and the lipidome. JCI Insight. 2018;3(12).

33. Turner SM, Linfoot PA, Neese RA, Hellerstein MK. Sources of plasma glucose and liver glycogen in fasted ob/ob mice. Acta Diabetol. 2005;42(4):187–93.

34. Bajaj M, Suraamornkul S, Pratipanawatr T, Hardies LJ, Pratipanawatr W, Glass L, et al. Pioglitazone reduces hepatic fat content and augments splanchnic glucose uptake in patients with type 2 diabetes. Diabetes. 2003;52(6):1364–70.

35. Raichur S, Wang ST, Chan PW, Li Y, Ching J, Chaurasia B, et al. CerS2 haploinsufficiency inhibits β-oxidation and confers susceptibility to diet-induced steatohepatitis and insulin resistance. Cell Metab. 2014;20(4):687–95.

36. He Q, Han X. Cardiolipin remodeling in diabetic heart. Chem Phys Lipids. 2014;179:75–81.

37. Cappel DA, Deja S, Duarte JAG, Kucejova B, Inigo M, Fletcher JA, et al. Pyruvate-Carboxylase-Mediated Anaplerosis Promotes Antioxidant Capacity by Sustaining TCA Cycle and Redox Metabolism in Liver. Cell Metab. 2019;29(6):1291–305 e8.

38. Lee MH, DeBerardinis RJ, Wen X, Corbin IR, Sherry AD, Malloy CR, et al. Active pyruvate dehydrogenase and impaired gluconeogenesis in orthotopic hepatomas of rats. Metabolism. 2019;101:153993.

39. Huang B, Wu P, Bowker-Kinley MM, Harris RA. Regulation of pyruvate dehydrogenase kinase expression by peroxisome proliferator-activated receptor-alpha ligands, glucocorticoids, and insulin. Diabetes. 2002;51(2):276–83.

40. Wu P, Blair PV, Sato J, Jaskiewicz J, Popov KM, Harris RA. Starvation increases the amount of pyruvate dehydrogenase kinase in several mammalian tissues. Arch Biochem Biophys. 2000;381(1):1–7.

41. Divakaruni AS, Wiley SE, Rogers GW, Andreyev AY, Petrosyan S, Loviscach M, et al. Thiazolidinediones are acute, specific inhibitors of the mitochondrial pyruvate carrier. Proc Natl Acad Sci U S A. 2013;110(14):5422–7.

42. Perry RJ, Camporez JG, Kursawe R, Titchenell PM, Zhang D, Perry CJ, et al. Hepatic acetyl CoA links adipose tissue inflammation to hepatic insulin resistance and type 2 diabetes. Cell. 2015;160(4):745–58.

43. Puri P, Baillie RA, Wiest MM, Mirshahi F, Choudhury J, Cheung O, et al. A lipidomic analysis of nonalcoholic fatty liver disease. Hepatology. 2007;46(4):1081–90.

44. Cole LK, Mejia EM, Vandel M, Sparagna GC, Claypool SM, Dyck-Chan L, et al. Impaired Cardiolipin Biosynthesis Prevents Hepatic Steatosis and Diet-Induced Obesity. Diabetes. 2016;65(11):3289–300.

45. Han X, Yang J, Yang K, Zhao Z, Abendschein DR, Gross RW. Alterations in myocardial cardiolipin content and composition occur at the very earliest stages of diabetes: a shotgun lipidomics study. Biochemistry. 2007;46(21):6417–28.

46. Li J, Romestaing C, Han X, Li Y, Hao X, Wu Y, et al. Cardiolipin remodeling by ALCAT1 links oxidative stress and mitochondrial dysfunction to obesity. Cell Metab. 2010;12(2):154–65.

47. Fritz IB, Yue KT. LONG-CHAIN CARNITINE ACYLTRANSFERASE AND THE ROLE OF ACYLCARNITINE DERIVATIVES IN THE CATALYTIC INCREASE OF FATTY ACID OXIDATION INDUCED BY CARNITINE. J Lipid Res. 1963;4:279–88.

48. Pérez-Carreras M, Del Hoyo P, Martín MA, Rubio JC, Martín A, Castellano G, et al. Defective hepatic mitochondrial respiratory chain in patients with nonalcoholic steatohepatitis. Hepatology. 2003;38(4):999–1007.

49. Warshauer JT, Lopez X, Gordillo R, Hicks J, Holland WL, Anuwe E, et al. Effect of pioglitazone on plasma ceramides in adults with metabolic syndrome. Diabetes Metab Res Rev. 2015;31(7):734–44.

50. Srere PA. Citrate Synthase. Methods in Enzymology. 1969;13:3–11.

51. Han X, Yang K, Yang J, Cheng H, Gross RW. Shotgun lipidomics of cardiolipin molecular species in lipid extracts of biological samples. J Lipid Res. 2006;47(4):864–79.

52. Wang C, Liu F, Frisch-Daiello JL, Martin S, Patterson TA, Gu Q, et al. Lipidomics reveals a systemic energy deficient state that precedes neurotoxicity in neonatal monkeys after sevoflurane exposure. Anal Chim Acta. 2018;1037:87–96.

53. Wang M, Wang C, Han X. Selection of internal standards for accurate quantification of complex lipid species in biological extracts by electrospray ionization mass spectrometry-What, how and why? Mass Spectrom Rev. 2017;36(6):693–714.

54. Yang K, Cheng H, Gross RW, Han X. Automated lipid identification and quantification by multidimensional mass spectrometry-based shotgun lipidomics. Anal Chem. 2009;81(11):4356–68.

55. Ramaswamy V, Hooker JW, Withers RS, Nast RE, Brey WW, Edison AS. Development of a ^13^C-optimized 1.5-mm high temperature superconducting NMR probe. J Magn Reson. 2013;235:58–65.

56. Merritt M, Bretthorst GL, Burgess SC, Sherry AD, Malloy CR. Sources of plasma glucose by automated Bayesian analysis of 2H NMR spectra. Magn Reson Med. 2003;50(4):659–63.

57. Constantin-Teodosiu D, Cederblad G, Hultman E. A sensitive radioisotopic assay of pyruvate dehydrogenase complex in human muscle tissue. Anal Biochem. 1991;198(2):347–51.

58. Harris RC, Hultman E, Nordesjö LO. Glycogen, glycolytic intermediates and high-energy phosphates determined in biopsy samples of musculus quadriceps femoris of man at rest. Methods and variance of values. Scand J Clin Lab Invest. 1974;33(2):109–20.

